# Human hippocampal neurons track moments in a sequence of events

**DOI:** 10.1101/2020.12.17.423193

**Authors:** Leila Reddy, Benedikt Zoefel, Jessy K. Possel, Judith C. Peters, Doris Dijksterhuis, Marlene Poncet, Elisabeth C.W. van Straaten, Johannes C. Baayen, Sander Idema, Matthew W. Self

## Abstract

An indispensable feature of episodic memory is our ability to temporally piece together different elements of an experience into a coherent memory. Hippocampal “time cells” – neurons that represent temporal information – may play a critical role in this process. While these cells have been repeatedly found in rodents, it is still unclear to what extent similar temporal selectivity exists in the human hippocampus. Here we show that temporal context modulates the firing activity of human hippocampal neurons during structured temporal experiences. We recorded neuronal activity in the human brain while patients learned predictable sequences of pictures. We report that human time cells fire at successive moments in this task. Furthermore, time cells also signaled inherently changing temporal contexts during empty 10-second gap periods between trials, while participants waited for the task to resume. Finally, population activity allowed for decoding temporal epoch identity, both during sequence learning and during the gap periods. These findings suggest that human hippocampal neurons could play an essential role in temporally organizing distinct moments of an experience in episodic memory.

**Significance Statement:** Episodic memory refers to our ability to remember the “what, where, and when” of a past experience. Representing time is an important component of this form of memory. Here, we show that neurons in the human hippocampus represent temporal information. This temporal signature was observed both when participants were actively engaged in a memory task, as well as during 10s-long gaps when they were asked to wait before performing the task. Furthermore, the activity of the population of hippocampal cells allowed for decoding one temporal epoch from another. These results suggest a robust representation of time in the human hippocampus.

## Introduction

Creating episodic memories requires linking together distinct events of an experience with temporal fidelity. The brain must represent the temporal flow and order of events, and glue them together in the correct sequential order. “Time cells” in the hippocampus and adjacent structures might play an essential role in this temporal organization of memory (Hasselmo, 2009; Eichenbaum, 2014; Howard et al., 2014). In rodents, time cells signal changing temporal contexts in a variety of paradigms (Manns et al., 2007; Pastalkova et al., 2008; MacDonald et al., 2011; Kraus et al., 2013; MacDonald et al., 2013; Kraus et al., 2015). They fire at successive moments of time during a fixed interval and the activity of the population of time cells covers the entire time interval (Pastalkova et al., 2008). More recently, another class of “ramping cells” in the lateral entorhinal cortex has been discovered. Ramping cells show slowly rising or decaying activity with time, over a range of time scales. Temporal epoch identity could be decoded from the firing activity of the population of cells (Tsao et al., 2018).

Temporal coding has also been observed in neuronal activity patterns in the human hippocampus. For instance, neuronal activity in the human medial temporal lobe shows gradual changes over time in memory tasks (Howard et al., 2012; Folkerts et al., 2018). The recall of a particular item is accompanied by the reinstatement of its initial temporal representation (Gelbard-Sagiv et al., 2008; Howard et al., 2012; Folkerts et al., 2018). More recently, single neurons have also been shown to be modulated by time, akin to time cells in rodents, during encoding and retrieval in a free recall memory task (Umbach et al., 2020).

In the current study, we ask if human hippocampal neurons represent temporal information during sequential order learning. A large body of work in animals and humans has shown that the hippocampus is essential for remembering the temporal order of sequential events (Eichenbaum, 2013). For example, in humans, the hippocampus is activated when subjects recall the order of objects, and conversely, patients with hippocampal damage have trouble in temporal order judgements (Spiers et al., 2001; Ekstrom and Bookheimer, 2007). In animals, rats with hippocampal damage are impaired at remembering the sequential order of odors (Fortin et al., 2002). Given the importance of the hippocampus in sequence order learning and temporal order judgements, we tested whether human hippocampal neurons represented temporal information while participants learned the order of a sequence of items. We tested for temporal modulation of hippocampal activity in two experiments: (1) during sequence learning (Figure 1A), and (2) during empty gap periods inserted in the task during which participants passively waited for the sequence to resume (Figure 1B). Note that in these gap periods, any potential temporal information is not driven by external stimuli or events, but rather represents inherent changes in the patients’ moment-to-moment experience. We report that human hippocampal neurons fire at successive moments during these structured time periods, both while subjects actively monitor a sequence, as well as during empty temporal gaps between events.

**Figure 1.**
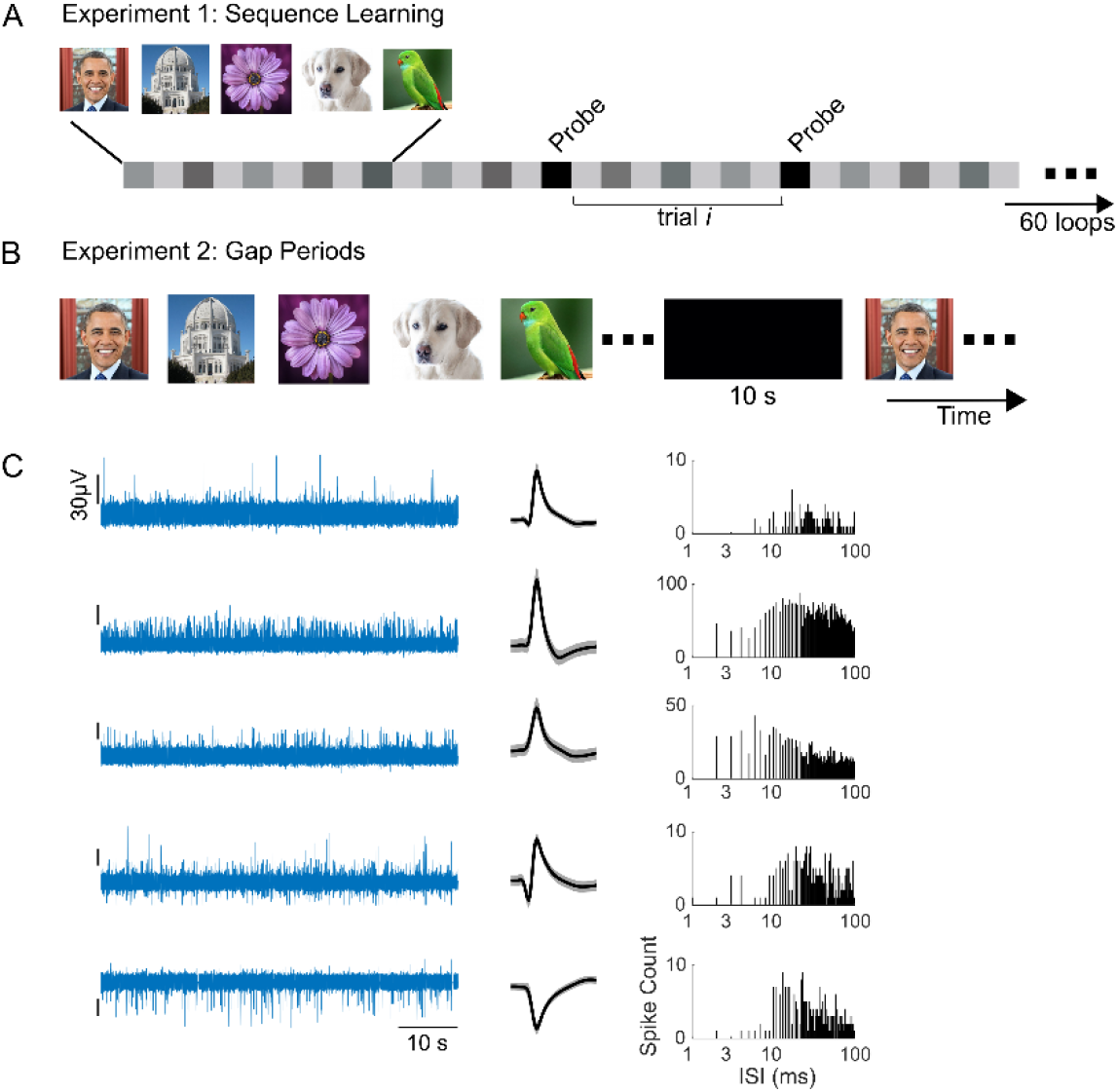
A, B) Experimental Design. In the sequence learning experiments, participants saw a sequence of images in a fixed order, and were asked to learn the sequence order. The stimulus sequence consisted of 5-7 image periods (image number fixed per session and determined by the availability of the patient) separated by inter-stimulus interval (ISI) periods. Each image was presented for 1.5s followed by an ISI of 0.5s. The sequence was repeated for 60 loops. 20% of the time, a probe event occurred (black squares) during which participants had to decide which of two choice images was the correct one at the current position in the sequence. The probe events occurred at random positions of the sequence. After the probe event, the sequence resumed. In our main analysis, we consider time periods that occurred between two consecutive probe events as the “trials” of interest. As shown, each post-probe “trial” consisted of several image and ISI periods (gray squares). B) The design of Experiment 2 was similar to that of Experiment 1 except for the insertion of 10-second-long gap periods (black rectangle) during sequence learning. These gap periods occurred periodically (see Methods). During the gap periods, the sequence stopped and patients were presented with a blank screen. They were asked to simply wait until the sequence resumed. C) Band-pass filtered (300-3000Hz) signal from five different channels (left), mean waveforms recorded on these channels (middle), and the corresponding distributions of inter-spike intervals (right). The black vertical tick marks on the left plots indicate a scale of 30uV.

## Materials and Methods

In this study, human epileptic patients performed two sequence learning tasks, while single neuron activity was recorded from microelectrodes implanted in the hippocampus. We quantified the influence of time on the firing activity of individual neurons using a stepwise general linear model (GLM), as has previously been used in the rodent literature (MacDonald et al., 2011; Tsao et al., 2018). In this GLM, a predictor variable is included in the model only if it is found to significantly improve the prediction of the response variable (see below).

### Patients

Nine patients of either sex participated in the first experiment, and six patients of either sex participated in the second experiment. The patients were diagnosed with pharmacologically intractable epilepsy, and were undergoing epileptological evaluation at the Amsterdam University Medical Center, location VUmc, The Netherlands. Patients were implanted with chronic depth electrodes for 7-10 days in order to localize the seizure focus for possible surgical resection (Fried et al., 1997; Engel et al., 2005). All surgeries were performed by J.C.B and S.I. The Medical Ethics Committee at the VU Medical Center approved the studies. The electrode locations were based entirely on clinical criteria and were evaluated based on the pre-surgical planned trajectories on the basis of structural MRI scans. The accuracy of the implantation was always checked using a CT scan co-registered to the MRI. We only included electrodes that were within a 3mm deviation from the target (based on visual confirmation). Each electrode contained eight microwires (Behnke-Fried electrodes, Ad-Tech Medical) from which we recorded multi-unit activity, and a ninth microwire that served as a local reference. The signal from the microwires was recorded using a 64-channel Neuralynx system, filtered between 1 and 9000 Hz, sampled at 32KHz. On average, each patient was implanted with 34 ± 11.8 microwires (range = [16, 48]). Participants sat in their hospital room at the Epilepsy Monitoring Unit, and performed the experimental sessions on a laptop computer.

### Spike Detection and Sorting

Spike detection and sorting were performed with wave_clus (Quiroga et al., 2004). Briefly (see (Reddy et al., 2015) for details), the data were band pass filtered between 300-3000Hz and spikes were detected with an automatic amplitude threshold. Spike sorting was performed with a wavelet transform that extracted the relevant features of the spike waveform. Clustering was performed using a super-paramagnetic clustering algorithm. Clusters were visually reviewed by the first-author for 1) the mean spike shape and its variance; 2) the ratio between the spike peak value and the noise level; 3) the inter spike interval distribution of each cluster; 4) the presence of a refractory period; 5) the similarity of each cluster to other clusters from the same microwire. Based on manual inspection of these criteria, clusters were retained, merged or discarded.

### Experimental design and statistical analyses

#### Behavioral Task

Experiment 1: Sequence Learning: The patients performed a total of 31 sequence learning (SL) sessions (Figure 1A). In each SL session, participants were presented with a sequence of 5-7 images (image number determined as a function of the difficulty level and the availability of the patient). The images were always presented in a pre-determined order such that a given image, A, predicted the identity of the next image, B, and so on. Subjects were asked to remember the order of the images in the sequence. Each image was presented for 1.5s (“image period”) followed by an “inter-stimulus interval period” (ISI) of 0.5s. The sequence was repeated continually 60 times resulting in experimental sessions of 10 minutes for 5-image sequences and 14 minutes for 7-image sequences, not including time spent by the subject to respond on probe events. On a random 20% of image periods, the sequence stopped and participants were presented with probe events. In these probe events, instead of being presented with the next image of the sequence, subjects were shown two images side by side and asked to decide (by pressing one of two keys on the keyboard) which of the two was the next image in the sequence. After the subjects had responded, the sequence resumed.

From the point of view of the subject, the probe events were salient moments of an otherwise repetitive experiment because the probes stopped the sequence and tested subjects on their learning of the sequence order. Thus, we considered sequence segments between two consecutive probe events as our “trials” of interest: structured, temporal experiences between two salient markers. We asked whether hippocampal neurons tracked time in this fixed interval. In control analyses described below, we verified these results with respect to other time periods in the experiment.

Experiment 2: Sequence learning with temporal gaps (Figure 1B): Six new patients performed eight sessions of a second SL experiment. This second experiment followed the design of the first SL experiment described above, except for the following modifications: 1) After a fixed number of repeats of the sequence, a 10s-long empty gap interval was presented. During these gap intervals, participants were presented with a black screen, without any stimulus input. They were asked to simply wait until the sequence started again. For three participants these gap intervals occurred after every 5 repeats of the sequence (resulting in 6 gap intervals in the experiment), while for the remaining three participants these gap intervals occurred after every 2 repeats of the sequence (resulting in 15 gap intervals in the experiment). 2) The sequence was repeated only 30 times instead of 60 times.

In the nine patients who performed the first experiment, we recorded from 441 neurons in the hippocampus, and in the six patients who performed the second experiment, we recorded 96 hippocampal units.

### Time Cell Identification with a General Linear Model (Experiment 1)

Time cell identification was performed with a GLM as in previous studies (MacDonald et al., 2011; Tsao et al., 2018). The firing activity of each neuron was modelled as a function of time, image identity, and whether the temporal period corresponded to an image or ISI period.

For the purposes of the GLM, as described above (Figure 1A), we defined “trials” as segments of the sequence between two consecutive probe events (number of sequence segments or “trials” between two consecutive probe events across sessions: mean ± s.e.m= 73.6±2.4). We made this choice because (i) as explained previously, these probe events were the most salient events of the experiment, and (ii) if we simply consider time=0 as the start of each 5-7 image sequence, “time” in the sequence is directly confounded by image identity because the sequence order is fixed. By redefining time=0 as the time at which the sequence restarted after the probe events, we avoided this confound because time is not confounded with image identity with respect to probe events (the sequence segment after the probe is random since the probe events occurred at random moments). In control analyses we also considered different temporal intervals for determining time cells.

Each of the post-probe “trials” consisted of several image and ISI periods that regularly followed each other (Figure 1A). The median number of image and ISI periods in a “trial” was seven, corresponding to a median trial length of 6.5 seconds. Thus, each “trial” between probe events was a well-structured temporal interval during which the sequence progressed according to its fixed order.

In the GLM, the firing activity vector (*Y*) on each “trial” contained the average firing rates for each period of the trial, with no smoothing or additional preprocessing. *Y* was modelled as a function of three variables: image identity in each period, time of the period (i.e., time of the mid-point of the period with respect to trial onset/probe offset), and whether the period corresponded to an image or an ISI event. A linear factor for time assumes that time cells either show a ramping up or a ramping down of firing activity during the trial. To also include the possibility of cells having a preferred time not just at the beginning and end of trials, but also in the middle of trials, we included a quadratic time term (i.e., *t*^2^, a parabola-shaped function identifying cells that would have stronger or weaker activity in the middle of the time window; for this purpose, time was recentered to the middle of the trial so that the parabola was centered in the middle of the time window).

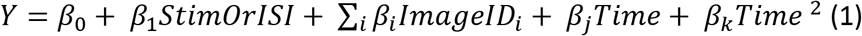

The GLM analysis was performed using the MATLAB *stepwiselm* function, including the variables *stim/ISI, imageID, Time* and *Time*^2^, in a linear model, with a constant term as the baseline model, the SSE criterion *(PEnter* = 0.05), and other default parameters. The variables *stim/ISI* and *imageID* were entered as categorical variables, and the time variables were continuous variables. Stepwise regression systematically tests the variance explained by adding and removing variables from a linear model based on their statistical significance in explaining the response variable. Note that the order in which regressors are entered into the stepwise linear model does not affect its outcome. Time cells were defined as cells for which the time terms (i.e., *Time* and/or *Time^2^)* were added by the stepwiselm function *(PEnter* < 0.05).

As a separate test to confirm our classification of time cells, different from the stepwise regression test, we performed a likelihood ratio test to compare the log-likelihood values of a restricted linear model which included all terms except the time terms, and a full model which also included the two time terms.

### Time Cell Identification with a General Linear Model (Experiment 2)

The 10s gap intervals of experiment 2 were epoched into 500ms non-overlapping windows and, as above, a stepwise regression analysis was performed: the firing activity *(Y)* in each epoch was modelled as a function of time in the epoch *(Time* and a quadratic time term *Time*^2^). All other parameters in this analysis were identical to those described for Experiment 1.

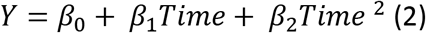

### Control analyses for defining trial periods and time cells

In the main analysis of Experiment 1, a “trial” was defined with respect to the probe events (i.e., the sequence segment that occurred between two consecutive probe events). We performed several additional analyses for identifying time cells. 1) Time cells were identified when the first period after the probe event was excluded from the GLM. 2) Time cells were identified when the GLM analysis was performed on only the ISI periods. In this control, the *Y* vector contained the firing rates in the ISI periods, and the regressor matrix *X* consisted of a time factor (time at the midpoint of the periods, and a quadratic term *Time*^2^) and an image identity factor (the identity of the image following the ISI period, to account for image-specific anticipatory responses that can be observed in the ISI periods during sequence learning (Reddy et al., 2015)). 3) “Trials” were re-defined as sequence segments with respect to the onset of each repetition of the sequence. Note that in this case, time selectivity can be confounded by stimulus selectivity (since the same stimulus sequence repeats identically in every loop); however, the Matlab *stepwiselm* function that we used for determining time selectivity could disentangle the potential contributions of the time and stimulus ID variables since it systematically tests for the addition and removal of each variable in significantly explaining the response variable. Nonetheless, to avoid any ambiguity in interpretation, we elected to present time selectivity with respect to probe events as our main analysis, as it precludes this potential confound.

### Statistical test for the number of time cells

Statistical significance for the number of time cells identified in the GLM analysis was evaluated using a permutation test. For the permutation test, in Experiment 1, surrogate data was created by randomly shuffling the image and ISI periods on each trial. For Experiment 2 surrogate data was created by randomly shuffling epoch time. The stepwise regression was then performed on this surrogate data, on 10^6^ iterations. The proportion of surrogates which had a higher number of time cells was *p* < 10^-6^ in all analyses (in other words, none of the surrogates ever had a higher number of time cells).

### Heatmaps cross-validation and statistical test

The “original” heatmaps shown in Figures 2B and 4B were constructed by sorting the time cells according to the latency of peak firing and using this sorted order to plot the firing rate of each cell over the time interval. However, a heatmap generated from random data, sorted and plotted according to the maximum value of each entry, will also show a similar well-organized pattern. Thus, to ensure the reliability of these maps, we performed a cross-validation analysis on the real heatmaps to verify their reproducibility.

**Figure 2.**
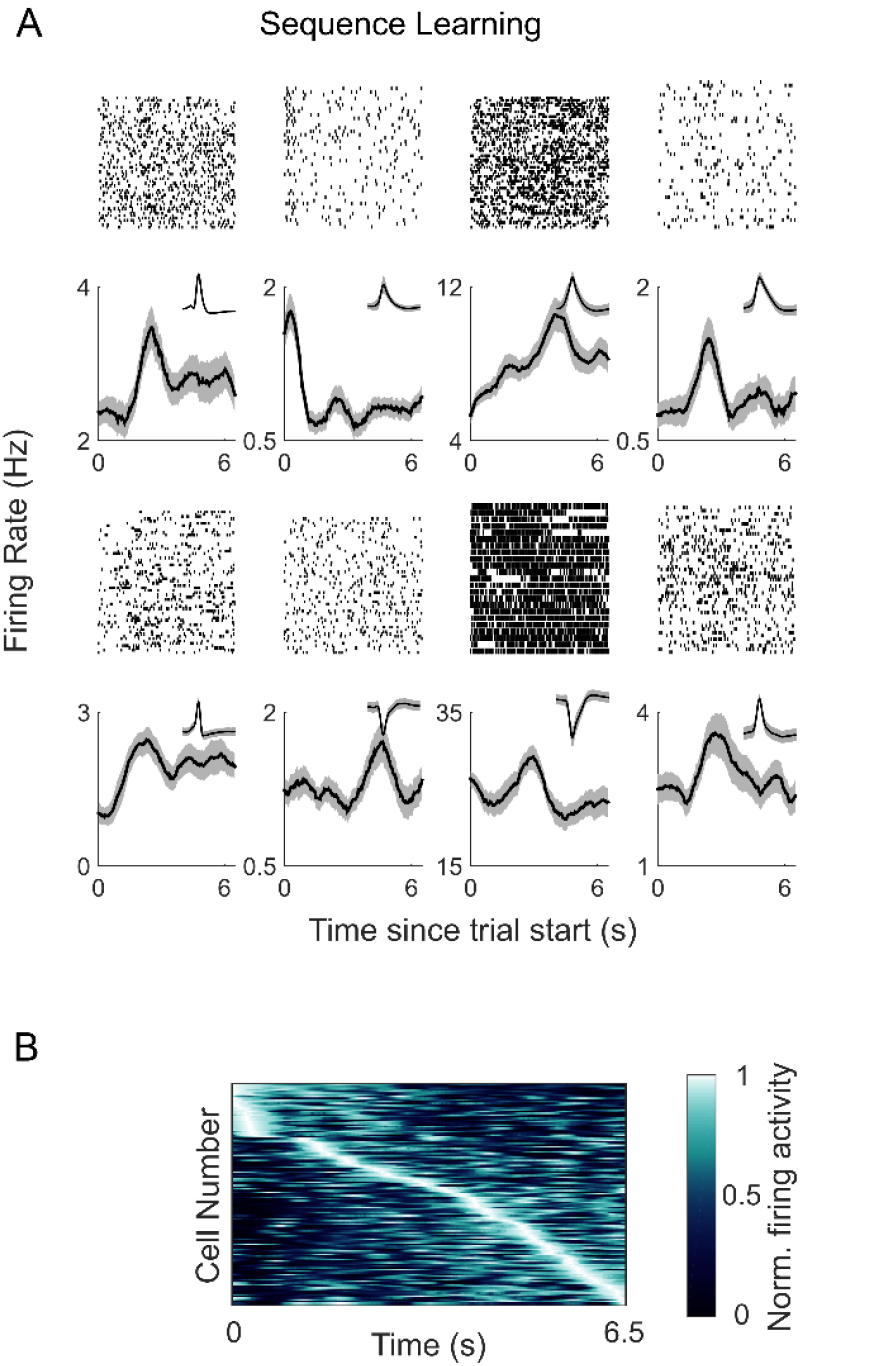
A) Hippocampal neurons are modulated by time during sequence learning. Raster plots (top) and post-stimulus time histograms (bottom) are shown for 8 example time cells. The x-axis corresponds to time of the median trial length (6.5s, see Methods). The black line is the average firing activity and the shaded area corresponds to the standard error of the mean across trials. Insets show the waveforms for these cells. B) Hippocampal neurons fire at successive moments of a temporal interval. Firing activity of the population of time cells (*N*=132) identified as being significantly modulated by time (i.e., ‘time’ and/or ‘time^2^’) in the sequence learning experiment. Each row shows the firing activity for an individual time cell, averaged across trials. The x-axis corresponds to time of the median trial length. The neurons are sorted by the latency of the maximum firing rate.

For the cross-validated heatmaps, the order in which the cells were plotted was determined from the latencies on a random half of the data and the firing rates were plotted for the remaining half of the data. If the cells had no true temporal preference, the peak latency on the second half should be unrelated to the latency measured on the first half, and no meaningful ordering of cells should appear. This cross-validation procedure was repeated 1000 times, and the resulting heat maps were averaged across cross-validations to generate cross-validated maps. We quantified the reliability of the heatmaps by performing a non-parametric permutation test. We first computed the Spearman correlation point-wise between the average cross-validated heatmap and the original heatmap, resulting in a correlation measure corr_orig,cv_. To simulate the null hypothesis that the cells do not have a reliable time preference, on each cross-validation iteration we generated 10^5^ surrogate heatmaps by randomizing the cell order obtained from the first half of the data (instead of determining their order based on peak firing time over the first half), and generating a heatmap with this random order for the second half of the data. Each of these 10^5^ surrogate heatmaps was correlated with the original heatmap (corr_orig,surr_). After averaging across the 1000 cross-validations, none of the 10^5^ surrogates yielded a correlation value corr_orig,surr_ higher than the real correlation corrorig,cv, i.e., *p*<10^-5^.

### Population pattern analysis

For this analysis, the population size was 441 neurons for Experiment 1, and 96 neurons for Experiment2.

Experiment 1: Sequence learning: The population pattern analysis for Experiment 1 was performed on the image periods. As described above, each “trial” (i.e., the sequence segment between two consecutive probe events) consisted of a varying number of image periods (Figure 1A). In the population pattern analysis our goal was to discriminate temporal period identity (e.g., image period 1 vs. image period 2 etc.). Discrimination performance was evaluated for different numbers of temporal periods from the start of the trial (ranging from two to five; i.e., discriminating between the first two image periods after the probe event, the first three image periods and so on). To ensure that performance was not driven by an unequal representation of the different images in the different periods, we created a balanced dataset for the population pattern analysis. In this balanced dataset we included the subset of trials per cell that assured that each image was equally present in each period (Figure 5A). The balanced dataset only included cells for which a minimum number of trials had been recorded. The minimum number of trials was the smallest number such that at least 300 cells were included in the analysis.

For the 2-way discrimination (period 1 vs. period 2), the balanced dataset required 42 trials, and 397 cells were included; for a 3-way discrimination (period 1 vs. period 2 vs. period 3) it required 30 trials, and included 303 cells; for a 4-way discrimination it required 12 trials and included 320 cells; and for a 5-way discrimination, it required 6 trials and included 367 cells. Discrimination of temporal period identity in this balanced dataset was thus not influenced by an imbalance in the number of presentations of each image across temporal periods. Note that the balanced dataset did not include the first ISI period after the probe and decoding was thus not influenced by the offset of the probe events. As mentioned below, trial selection for creating the balanced dataset was repeated on 100 iterations, and the population pattern analysis was performed on each iteration.

The population pattern analysis was performed using a split-half approach (Haxby et al., 2001) on the firing rates of the balanced dataset (Figure 5B-D). The trials in the balanced dataset were randomly split into two halves, and in each half the firing activity of the population of neurons was arranged into vectors, per period and per trial (Figure 5B). These vectors were averaged across trials, yielding a population pattern vector for each period (“period vectors”) in each half of the dataset (Figure 5C). To quantify decoding or discrimination performance, we pairwise correlated the period vectors in one half of the dataset with the period vectors in the other half, and used the pairwise correlation values to measure the percentage of correct classification (Figure 5D). To be more precise, for each period we computed the correlation distance for this period across the two halves of the dataset (“within” comparison), and compared it to the correlation distance with a different period in the other half of the dataset (“between” comparison). Decoding was correct if the distance for the “within” comparison was lower than the distance for the “between” comparison (Haxby et al., 2001). This procedure was repeated for all pairs of periods, and decoding accuracy was the proportion of correct comparisons. Feature normalization was performed on the dataset for this analysis by performing a z-score on the data for each cell along the periods dimension. Feature normalization was performed on the whole dataset based on the mean and standard deviation measured in the “training” half of the dataset (to avoid “leakage” of information from the training half to the test half during the split-half crossvalidation).

Note that in the split-half approach the “training data” (one half of the data) and “testing data” (the other half of the data) are independent by construction.

To increase reliability, the population pattern analysis was performed over several iterations: (i) trial selection for the balanced dataset was repeated 50 times; (ii) on each of these 50 iterations, the dataset was randomly split into two halves 200 times. The reported mean decoding accuracy was the average decoding performance across the 50 iterations for creating the balanced dataset. The standard deviation was computed across the 50 balanced datasets of the mean decoding accuracy across the 200 split halves. Statistical significance was computed with a t-test against chance (0.5) across the 50 iterations for creating the balanced dataset (all *p* < 10^-3^).

Experiment 2: Sequence learning with temporal gaps: For the population pattern analysis of the gap intervals, each gap interval was split into four, five or ten discrete periods and discrimination of temporal period identity was performed using a split-half approach (50 iterations). The procedure was the same as the one described above for Experiment 1 (Figure 5). Feature normalization was performed on the dataset by performing a z-score on the data. Statistical significance was tested using a t-test against chance performance (0.5) across the 50 split-half iterations. Since some subjects had 6 gap periods while others had 15 gap periods, on each of the 50 iterations for splitting the data a random set of 6 gap periods was chosen from the datasets which contained 15 gap periods.

## Results

### Behavioral task and number of units

To determine whether human hippocampal neurons are modulated by temporal context, we recorded from hippocampal neurons in human patients who performed a sequence learning task (Figure 1 A, B). We performed two independent sequence learning experiments in two groups of patients implanted with intracranial micro-electrodes. In Experiment 1, we recorded the activity of 441 hippocampal neurons in nine patients, and in Experiment 2 we recorded from 96 hippocampal neurons in a new group of six patients.

In both experiments, the patients were presented with a fixed number of images (5-7, depending on the patients’ availability) in a pre-defined order (Figure 1 A, B), and asked to learn the sequence order (Reddy et al., 2015). The sequence was repeated continually 60 times. Experiment 2 was similar to Experiment 1 except for the periodic insertion of 10-second-long gap periods during the experiment. During these gap periods, the sequence stopped and the patients had to wait until the sequence resumed (Figure 1B).

The duration of each image period in the sequence was 1.5s, and each image period was followed by an inter-stimulus interval (ISI) of 0.5s. On a random 20% of image periods, subjects were probed on their learning of the image order. During these probe events, the sequence momentarily stopped and subjects were presented with two images from the sequence. Their task was to report which of the two images was the correct one at the current sequence position. The sequence then resumed until the next probe event. Participants rapidly learned the sequence order and achieved >90% performance on probe trials within the first six sequence presentations (Reddy et al., 2015). From the point of view of the subjects the probe events were salient moments of the experiment because they stopped the ongoing sequence and tested learning.

### Human hippocampal neurons are modulated by time during sequence learning

Time cells have been characterized as neurons whose activity is modulated by temporal context within a well-defined time window. Our experiment design lent itself to identifying time cells because the task consisted of a structured image sequence that occurred in a fixed and predictable time interval. In the time domain, our experiment involved three distinct timelines: (1) experiment time running from the beginning to the end of the experiment, (2) sequence time with respect to the start of each iteration of the sequence, and (3) probe time running between consecutive probe events. We first focus on probe time since, as described above, the probe events were salient moments from the point of view of the participants. Furthermore, focusing on probe time allowed us to decouple time from image identity, since the post-probe trials consisted of varying segments of image and ISI periods.

We defined “trials” as segments of the experiment that occurred between two consecutive probe events in Experiment 1 (Figure 1A). These trials were of a relatively fixed duration (median trial length = 6.5 seconds), and consisted of a sequence of image and ISI periods. To identify time cells, we examined whether the firing activity of hippocampal neurons was modulated by time. Previous studies have identified time cells using a variety of frameworks such as fitting a Gaussian function to firing activity (Park et al., 2014; Salz et al., 2016), a one-way ANOVA of time and firing rate (Umbach et al., 2020), or a general linear model (GLM) factoring in the influence of time and other experimental factors on firing activity (MacDonald et al., 2011; Tsao et al., 2018). In the current study we elected to use the stepwise general linear model (GLM) method established by Tsao and colleagues (Tsao et al., 2018), since it allows us to identify time cells while also measuring the influence of the other experimental parameters on hippocampal responses (see the section *Human hippocampal neurons are “multidimensional”* below).

In the stepwise GLM framework used here, a predictor variable is included in the model only if it is found to significantly improve the prediction of the response variable (firing rate). For each neuron, we modelled the firing rate in each image/ISI period of the trial sequence with different potential predictors: image identity, period type (i.e., image or ISI period), and two time terms. A linear time term was included in the model to identify cells that show a ramping up or ramping down of firing activity. A second quadratic term was included to allow for the possibility of cells having a preferred time not just at the beginning and end of trials, but also in the middle. Time cells were identified as those in which one or both of the time terms was selected for inclusion in the stepwise linear model. Statistical significance was evaluated using a permutation test in which the image and ISI periods on each trial were randomly shuffled on 10^6^ iterations, and the stepwise GLM was performed on each iteration.

We identified a significant number of hippocampal neurons (30%) that were modulated by time during sequence learning (132 of 441 neurons; this number was more than expected by chance based on a permutation test, *p*<10^-6^). Of these, 84 were modulated by the linear time term, 40 by the quadratic time term, and 8 by both time terms. Individual examples of time cells are shown in Figure 2A.

In a separate analysis to confirm our classification of neurons as time cells, we performed a likelihood ratio test to compare the log-likelihood values of a restricted linear model which included all terms except the time terms, and a full model which also included the time terms. 96% of the cells classified as time cells based on the stepwise regression method had a significantly higher log-likelihood value according to this likelihood ratio test (*p*<0.05). Conversely, only 2% of the cells that were not labelled as time cells based on stepwise regression were selected by the likelihood ratio test.

We conducted a number of control analyses to verify that the temporal modulation of firing rates was observed under different analysis parameters (Figure 3A). Time cells were identified when the GLM was performed on (1) the firing rates of only the ISI periods (87 cells, permutation test, *p*<10^-6^), and (2) when excluding the first ISI period after the probe (94 cells, permutation test, *p*<10^-6^). Time cells were also detected when “trials” were re-defined as sequence segments with respect to the onset of each repetition of the sequence (i.e., instead of with respect to the onset of the probe events, as in the main analysis). In this analysis, we identified 94 neurons that were significantly modulated by the time variables (permutation test, *p*<10^-6^). Note that in this case, time selectivity can be confounded by stimulus selectivity (since the same stimulus sequence repeats identically in every loop); however, the stepwise regression approach could disentangle the potential contributions of the time and stimulus identity variables since it systematically tests for the addition and removal of each variable in significantly explaining the response variable. Nonetheless, to avoid any ambiguity in interpretation, we have elected to present time selectivity with respect to *probe* events as our main analysis, as it precludes this potential confound.

**Figure 3.**
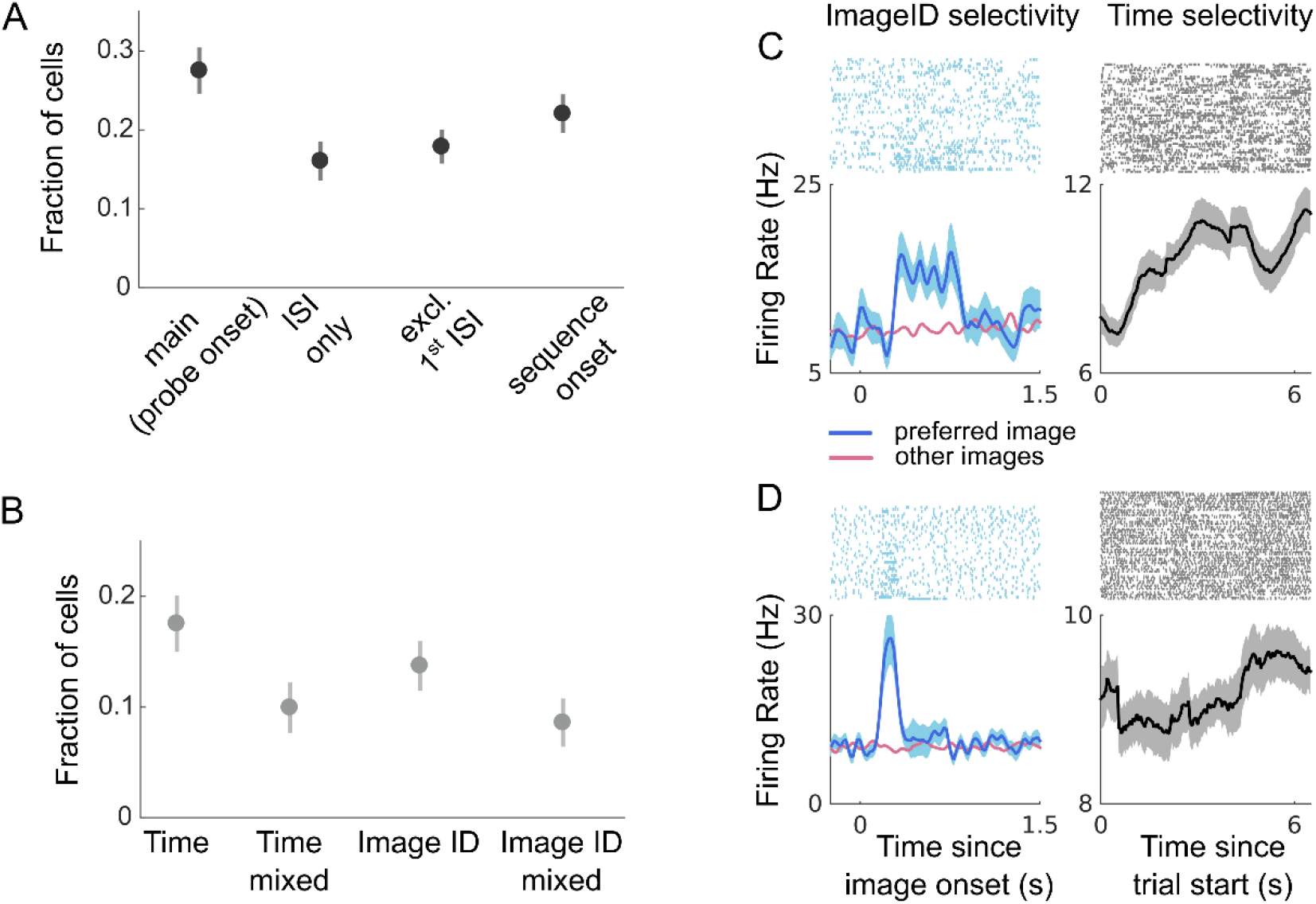
A) Fraction of cells that were modulated by time in the main analysis (i.e., with respect to the offset of the probe event), when time cells were identified on only the ISI periods (ISI only), when the first ISI period was excluded (excl. 1^st^ ISI), and when time cells were identified with respect to the onset of each iteration of the sequence (sequence onset). B) Hippocampal neurons are “multi-dimensional”. Fraction of cells that were exclusively modulated by time (Time), modulated by time and another variable (Time mixed, e.g. Time and ImageID), exclusively modulated by image type (ImageID), and modulated by image type and another variable (ImageID mixed). The dots and vertical lines correspond to the mean and standard error of the mean across recording sessions respectively (*N*=31 sessions). C, D). Examples of two cells that showed mixed selectivity for the factors of ImageID and time. *left).* These two cells were modulated by the factor of ImageID. The raster plots and post-stimulus time histograms are aligned to the onset of each image in the sequence. The blue curve is the response to the image in the sequence that elicited the highest response (preferred image). The red curve is the mean response to the other images in the sequence. The blue shaded area is the standard error of the mean across trials of image presentation. *right)* The same two cells were also modulated by the factor of Time. The raster plots and post-stimulus time histograms are now aligned to trial onset (i.e., the offset of the probe events). The format of the panels on the right is the same as Figure 2A. For visualization purposes only, and to better disentangle the effects of Time and ImageID, the spikes corresponding to the preferred images have been excised from the right panels.

Thus, human hippocampal neurons represent a changing temporal context while participants are actively engaged in memorizing the order of a sequence of events. Previous studies have shown that when considered at the population level, the firing activity of time cells covers the duration of a given temporal epoch (Pastalkova et al., 2008; MacDonald et al., 2011). Likewise, across all participants we observed neuronal peak firing at successive moments in time, and when each cell was ordered by its preferred moment of firing, population activity spanned the temporal window (Figure 2B). For consistency with previous studies, we illustrate this data as a population level heatmap. Although population level heatmaps are primarily used for display purposes, it can be informative to test their reliability, since random data sorted by peak value could also generate well-organized heat maps. To test the reliability of our heat maps we performed a cross-validation analysis, in which the preferred time of firing for each cell was determined in one half of the data and the consistency of time preference was measured in the second half of the data. Statistical significance of the cross-validated heatmaps was evaluated using a permutation method (Methods), in which cross-validated and surrogate (randomly permuted) heatmaps were correlated with the original heatmap. The proportion of surrogates which had a higher correlation than the cross-validated data was *p*<10^-5^, supporting the notion that temporal preference was reliable in our neuronal population.

### Human hippocampal neurons are “multidimensional”

In the rodent hippocampus, time cells are not exclusively modulated by time, just as place cells are not exclusively modulated by place. Rather, place cells and time cells appear to be “multidimensional” — they can be modulated by various factors, including the spatial, stimulus-related, and temporal dimensions of an experience (Eichenbaum, 2014).

In the human hippocampus, ~15% of neurons respond to visual stimuli (Quiroga et al., 2005). However, it is not yet known whether these neurons are exclusively visual, or if they are multidimensional and can be modulated by other factors, such as the temporal context of an experience. An advantage of the GLM-based approach used in the present study is that it allowed us to tease apart the influence of different experimental factors on the firing activity of hippocampal neurons (Figure 3B, C, D). The GLM analysis quantified the influence of stimulus presence (i.e., image periods vs. ISI periods), image identity, and time in the trial. A considerable number of neurons was modulated exclusively by time (92 out of 441, 17.5%) or image identity (56 out of 441, 13.7%), but we also found neurons selective for a combination of these factors (40 neurons, 9.9% for time and another factor; 30 neurons, 8.6% for image identity and another factor).

### Internally generated time selectivity in human hippocampal neurons

The results from Experiment 1 demonstrate that human hippocampal neurons are modulated by the temporal context of an explicit task. Do human hippocampal neurons also represent the temporal structure of an experience in the absence of external inputs or an overt task?

We performed a second experiment in six new patients who performed a different version of the sequence learning task (Figure 1B). In this new task, participants learned the sequence order as before, but every so often, the sequence stopped for 10s and participants waited until the sequence resumed. The participants had no stimulus input during these 10s gap periods -- they were presented with a blank screen. We isolated 96 hippocampal neurons in this second experiment, and used a GLM approach to determine whether human hippocampal neurons represent temporal information during the gap periods. As before, significance testing was based on a permutation test in which we repeated the GLM 10^6^ times after shuffling the data (Methods).

During the gap periods, 33 hippocampal neurons (34% of cells; Figure 4A) were significantly modulated by time (more than expected by chance based on a permutation test, *p*<10^-6^), whereas while the patients were engaged in sequence-learning 18 neurons (19% of cells) were time-selective. 8 neurons encoded temporal information during both the task period and the gap period, consistent with previous results showing that the recruitment of temporally sensitive cells can change with task demands or behavior, and that subsets of cells may overlap in different task contexts (Pastalkova et al., 2008; MacDonald et al., 2011; Tsao et al., 2018; Umbach et al., 2020). During the gap periods, the time of peak firing in the population occurred at successive moments, and population activity spanned the entire 10s interval (Figure 4B). As above, the reliability of the heat map was statistically verified with a cross-validation procedure (permutation test *p*<10^-5^, Methods). As reported in rodents, there was a stronger representation of earlier timepoints (Salz et al., 2016). Thus, even in the absence of visual input or an overt task, the firing activity of hippocampal neurons is inherently modulated by the changing temporal context.

**Figure 4.**
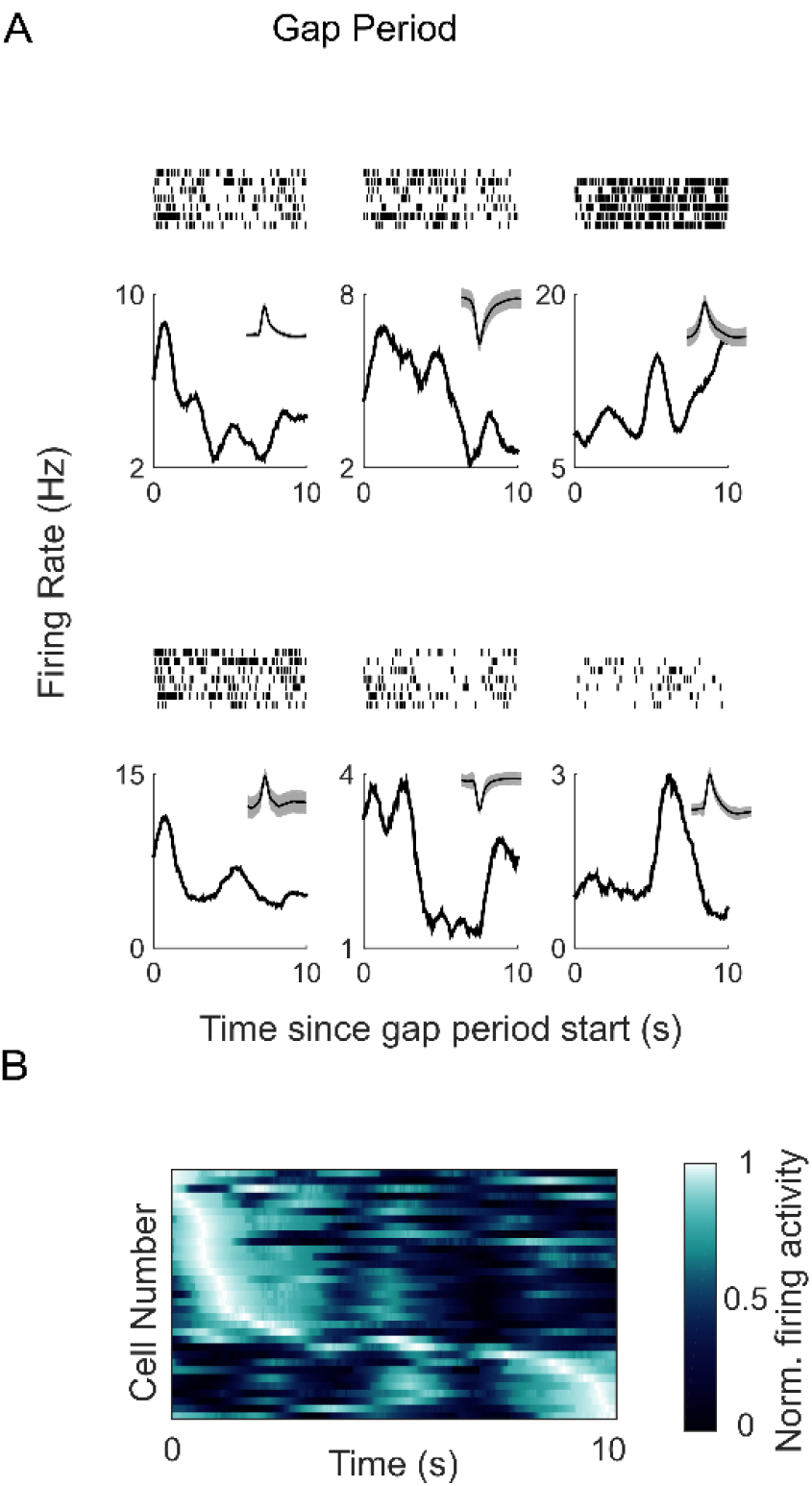
A) Hippocampal neurons are modulated by time during the gap periods. Raster plots (top) and post-stimulus time histograms (bottom) are shown for 6 example time cells. The x-axis corresponds to the 10s duration of the gap period. Insets show the waveform for these cells. B) Hippocampal neurons fire at successive moments of the gap period. Firing activity of time cells in the temporal gap experiment (*N*=33). Each row shows the firing activity for an individual cell, averaged across the gap periods. The x-axis corresponds to the 10s gap period. The neurons are sorted by the latency of the maximum firing rate.

### Hippocampal population activity encodes temporal information

In the rodent brain, time information is signaled both explicitly in individual neurons, and can also be gleaned from population-level dynamics of time-selective and non-time-selective cells (Tsao et al., 2018). Is time information also reflected in the population activity of human hippocampal neurons? To address this question, we performed a population pattern analysis of image period identity in the sequence learning sessions (Figure 5).

**Figure 5:**
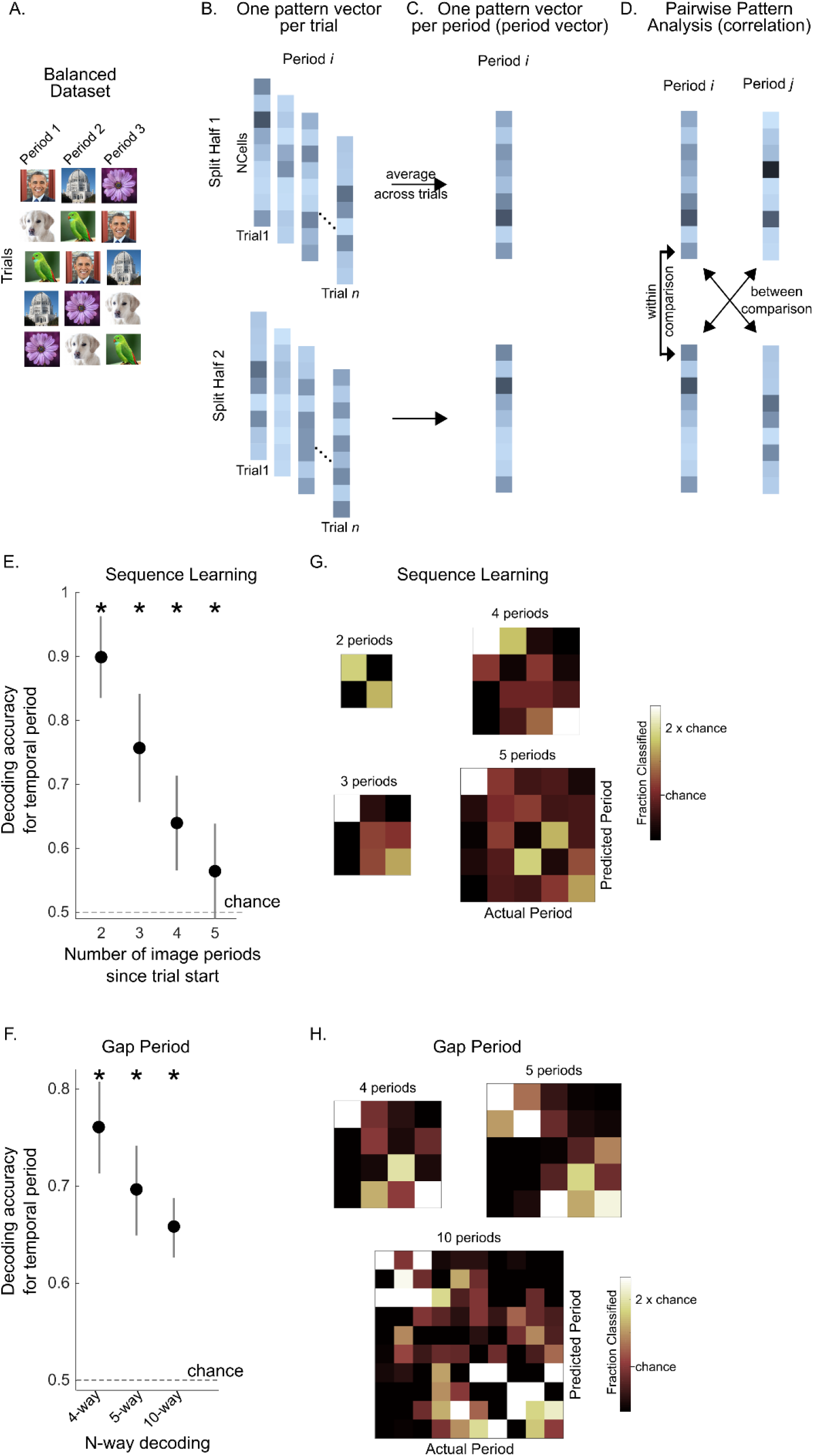
Population pattern analysis. In experiment 1, the population pattern analysis was performed on image periods (e.g., discriminating image period 1 vs image period 2). In experiment 2, the population pattern analysis was performed on the 10s gap interval. A). For experiment 1, a balanced dataset was created by selecting a subset of trials so that each image was equally present in each period across trials (see methods). B) Population pattern analysis procedure. The trials of the balanced dataset (experiment 1), or the gap intervals (experiment 2) were split into two halves (repeated on 100 iterations; see methods). In each half of the dataset, and for each period, the firing activity of the population of cells was arranged into a pattern vector for each trial. C). An average pattern vector of population firing activity was obtained for each image period by averaging across trials (period vectors). D). Pairwise discrimination was performed on the period vectors across the two halves of the dataset. For all pairs of periods, the Pearson’s correlation was computed across the two halves, and the same-period comparisons (“within comparisons”) were compared to the different-period (“between”) comparisons. Decoding was correct if the correlation for the “within” comparison was higher than the correlation for the “between” comparison (Haxby et al., 2001). E-H). Population pattern analysis accuracies. E). Pairwise decoding accuracy for temporal period identity during the sequence learning experiment using the split-half procedure described in A-D (population size=441 neurons). The x-axis shows the number of image periods that the classifier was tested on (i.e., discrimination between the first two image periods, the first three image periods, etc., since the start of the trial). Decoding accuracy (mean ± standard deviation) for discriminating the first two periods = 89.9±6.4%, of the first three periods = 75.7±8.5%, of the first four periods = 64.0±7.4%, of the first five periods = 55.7±8.2%. The black dots and error bars correspond to the mean and standard deviation of decoding performance across the 50 iterations for creating the balanced dataset. Asterisks denote significance based on a t-test against chance, *(p* < 10^-3^). F). Population pattern analysis decoding performance during the temporal gap experiment (population size=96 neurons). The 10s gap periods were split into four, five or ten discrete periods and discrimination of temporal period identity was computed using the procedure shown in B-D. Decoding performance was significantly above chance for all comparisons (t-test against chance (0.5), *p* < 10^-3^). Pairwise decoding accuracy, mean ± standard deviation = 76.0 ± 4.7% for 4-way decoding, 69.6 ± 4.6% for 5-way decoding, and 65.7 ± 3.1% for 10-way decoding. The black dots and error bars correspond to the mean and standard deviation of decoding performance across the 50 iterations of the split-half procedure. G, H) Decoding errors mainly occurred for predicting neighboring temporal periods. G) Confusion matrices during sequence learning when discriminating the first two temporal periods, the first three temporal periods etc. H) Confusion matrix for the gap experiment for different epoch lengths.

In Experiment 1, each trial of the experiment consisted of a sequence of image periods between two consecutive probes, and the goal of the population pattern analysis was to determine whether hippocampal population dynamics reflected the temporal identity of each image period (e.g., discriminate “image period 1” vs. “image period 2”). To ensure that decoding or discrimination performance was not driven by an unequal representation of the different images in the different image periods, we created a balanced dataset in which accurate decoding of the temporal identity of each image period cannot arise from merely decoding spurious image information (Figure 5A).

Discrimination performance was evaluated for different numbers of image periods from the start of the trial (ranging from two to five; i.e., discriminating between the first two image periods after the probe event, the first three image periods and so on). High decoding accuracy for temporal period identity was observed for discriminating all image periods (Figure 5E; mean ± standard deviation of decoding accuracy for discrimination of the first two periods = 89.9±6.4%, t(49) = 44.1, p < 0.001; of the first three periods = 75.7±8.5%, t(49) = 21.4, p < 0.001; of the first four periods = 64.0±7.4%, t(49) = 13.4, p < 0.001; of the first five periods = 55.79±8.2%, t(49) = 4.9, p < 0.001). Decoding errors occurred primarily for neighboring periods (Figure 5G). Decoding accuracy could not be biased by the offset of the probe events since the first ISI period after the probe event was excluded in the population pattern analysis. Thus, hippocampus population dynamics uniquely represented each temporal period.

Temporal epoch information was also present in population-level dynamics during the gap periods. A population pattern analysis during these gap periods revealed a significant representation of time information at the level of the overall population, and decoding errors mainly occurred for neighboring epochs (Figure 5F, H). High decoding accuracy for temporal epoch identity was observed for different temporal epoch sizes (mean accuracy ± standard deviation for 4-way decoding = 76.0 ± 4.7%, t(49) = 38.9, p < 0.001; for 5-way decoding = 69.6 ± 4.6%, t(49) = 29.9, p < 0.001; and for 10-way decoding = 65.4 ± 3.1%, t(49) = 36.3, p < 0.001). Decoding accuracy during the gap periods could not have been driven by external visual input or overt behavior; rather the high decoding accuracy reflects an internally generated temporal context signal represented in the population of neurons.

## Discussion

In this study we report that human hippocampal neurons represent temporal information as subjects progress through a sequence of events, and during empty gap periods in the sequence. Different neurons responded successively at different moments of the task, and together, the activity of these neurons covered the entire task period. This finding of time cells in the human hippocampus extends prior evidence for temporal coding reported in both rodents (Pastalkova et al., 2008; MacDonald et al., 2011) and humans (Folkerts et al., 2018; Umbach et al., 2020). Perhaps most relevant to the current study are the recent findings of Umbach and colleagues who showed time cell activity in the human hippocampus during a free recall memory task (Umbach et al., 2020). Similar to that study, we show that hippocampal neurons are modulated by time during sequence learning, providing further evidence in support of time cells in the human hippocampus.

Temporal information is represented at different time scales in the hippocampus and neighboring brain regions. In rodents, hippocampal time cells typically show sharp tuning for particular moments of a fixed interval (Pastalkova et al., 2008; MacDonald et al., 2011), whereas in the lateral entorhinal cortex, temporally sensitive neurons show a more gradual ramping of firing activity (Tsao et al., 2018). In humans, population activity gradually changes over a period of minutes (Folkerts et al., 2018), but single neurons can also show punctuated time-cell-like firing patterns as we show here and as previously reported (Umbach et al., 2020). The tuning of time cells in the human hippocampus appears to be broader than in rodents, both in the Umbach et al., study as well as in our study. Future work should test how the temporal precision of human time cells depends on task conditions, for example, in more difficult tasks that probe temporal order at finer timescales. Rodent time cells also display a scalar coding of time, such that cells that are active later in a time window, fire for longer periods. Further, just like place cells that remap, time cells have been observed to “re-time”: cells can change their temporal preferences within the same recording session when the temporal structure of the experience is changed (MacDonald et al., 2011). These observations in rodent time cells raise intriguing possibilities for future investigations of temporal coding in the human hippocampus.

The temporal modulation of time cells in our study was also observed during empty 10s gap periods in which patients were not presented with any visual input or required to perform an explicit task. Time cells were observed to fire at successive moments in these blank periods. Temporal modulation during these gap periods could not have been driven by external events; rather they appear to represent an evolving temporal signal as a result of changes in the patients’ experience during this time of waiting. Related to this point, temporal coding in the hippocampus and the lateral entorhinal cortex has been shown to change with behavior and task demands (MacDonald et al., 2013; Tsao et al., 2018). The design of Experiment 2 allowed us to directly examine whether cells that were modulated by time during sequence learning were also recruited during the gap periods when the patients did not overtly perform a task. We found that different subsets of cells were active during these two task periods, with a partial overlap of cells that were active in both periods. These results suggest a context-dependent recruitment of cells for the representation of temporal information.

In rodents, time cells and place cells do not uniquely represent temporal and spatial information respectively. Rather, medial temporal lobe neurons can be influenced by various experimental factors, including the stimulus-related, spatial and temporal facets of an experience (Komorowski et al., 2009; Tsao et al., 2018). Similarly, we found that human time cells were not exclusively modulated by time, but also encoded sensory information about the presence or absence of a stimulus, and the identity of the stimulus. Such multi-dimensional representations could play a critical role in episodic memory mechanisms in which the “what”, “where”, and “when” elements of an experience are bound together into a coherent memory.

The phenomenon of subjective “mental time travel” is a cornerstone of episodic memory (Tulving, 2002). Central to our experience of reliving the past is our ability to vividly recall specific events that occurred at a specific place and in a specific temporal order. Time cells in rodents and humans, and other temporally-sensitive populations of neurons support theoretical frameworks that posit that temporal context information plays an important role in memory mechanisms in the hippocampus (Howard et al., 2014; Howard et al., 2015). Our results provide further evidence that human hippocampal neurons represent the flow of time in an experience.

## Acknowledgements

This work was supported by grants from the French Agence Nationale de la Recherche (ANR-12-JSH2-0004-01), ANITI (Artificial and Natural Intelligence Toulouse Institute) Research Chair (ANR-19-PI3A-0004), the Fyssen foundation, and the Université Paul Sabatier, Toulouse, France (BQR, 2009 and Appel à Projets de Recherche Labellisés, 2013), to L.R. JP was supported by an NWO Onderzoekstalent grant awarded to MWS. DD was supported by an NWA-Startimpuls grant awarded to MWS. We would like to thank Pieter Roelfsema for his support.

